# The role of greater competitive ability in countering age disadvantages in larval competition in *Drosophila melanogaster*

**DOI:** 10.1101/2023.07.25.550469

**Authors:** Srikant Venkitachalam, Auroni Deep, Srijan Das, Amitabh Joshi

## Abstract

**Background:** Populations of *Drosophila melanogaster* adapted to high larval densities evolve increased larval competitive ability compared to low density controls. However, traits contributing to greater competitive ability can differ across such populations, depending on the exact details of high-density conditions imposed. In the current study, we consider three sets of *D. melanogaster* populations adapted to three different kinds of high-density scenarios. These population sets have evolved different degrees of increases in egg size and decreases in egg hatching time as correlates of increased larval competitive ability.

**Question:** We asked two related questions:

a. Do populations adapted to larval crowding counter an imposed initial age disadvantage in larval competition, better than their controls?
b. Do differences in egg size and egg hatching time among crowding-adapted populations matter in competitive performance when suffering age disadvantage?

**Study system:** We used three sets of outbred laboratory *Drosophila melanogaster* populations selected for larval crowding with different egg number, food volume and vial type combinations (named MCU, CCU and LCU). We also used one set of low-density ancestral control populations (named MB).

**Methods:** We implemented high density cultures with half the eggs from one of the study (focal) populations, and the other half from a common marked competitor population (Orange Eye: OE). We provided head start durations of 0, 3, 5, or 7 hours to the eggs of the common competitor. This imposed the respective age disadvantage to the larvae of the focal population. Pre-adult development time of survivors was the indicator of competitive performance used.

**Results:** All crowding-adapted populations countered an initial age disadvantage better than the control populations. We did not see any differences among crowding adapted populations regarding their performance in countering the age disadvantage. The common competitors showed the best competitive performance against the populations with the greatest egg size and shortest hatching time.

**Conclusions:** Adaptation to crowding leads to significantly better chances against age disadvantages in larval competition. Temporal head starts need not be of overwhelming advantage in larval competition against superior competitors in *Drosophila*. Among crowding adapted populations, evolving greater egg size and shorter hatching time does not seem to better offset the effects of age disadvantage to larvae. Differences in larval effectiveness and tolerance of the populations are likely to explain these results.

## Introduction

Competition holds a central place in the theory of density-dependent selection (MacArthur, 1962; MacArthur and Wilson, 1967), which is one of the few interfaces between ecology and evolution (reviewed in Mueller, 1997, 2009). A fundamental prediction of this theory is that a population adapted to crowded conditions will evolve greater average competitive ability (Mueller, 1988a). We define competitive ability as the sum of traits relevant to the individual’s performance in competition (Bakker, 1961): essentially, fitness under competitive conditions.

The traits contributing to competitive ability can be teased apart by studying laboratory populations adapted to high density conditions. Such long-term experiments would be expected to lead to the evolution of greater competitive ability over generations of selection. Over the past four decades, experimenters have used populations of *Drosophila* extensively for such selection experiments. The results from several selection experiments have explored the evolution of increased larval competitive ability and its contributing traits (reviewed in Mueller, 1997, 2009; Prasad and Joshi, 2003; Nagarajan *et al*., 2016; Venkitachalam *et al*., 2022).

However, over the decades, different selection experiments have shown very contrasting results regarding which traits underlie the greater competitive ability (Nagarajan *et al*., 2016; Sarangi *et al*., 2016; Sarangi, 2018). Due to this, we do not yet have a comprehensive understanding of the relative importance of different traits in the context of larval competition.

The early selection studies in the 1980s and 1990s did show some consistent results among themselves and led to a canonical understanding of which traits typically mediate the evolution of larval competitive ability in *Drosophila*. The first studies were done on *D*. *melanogaster* populations selected for high population densities at all life-stages (the *K*-populations), compared to their low population density controls (the *r*-populations) (Mueller and Ayala, 1981). The selection experiment met the predictions made by models of density-dependent selection. The *K*-populations evolved increased larval competitive ability compared to the *r*-populations (Mueller, 1988b). The *K*-population larvae also evolved increased larval feeding rate as well as increased larval foraging path length (Joshi and Mueller, 1988; Sokolowski *et al*., 1997). Both these traits could be essential to larval competition. However, the *K*-populations showed reduced food to biomass efficiency (Mueller, 1990), which was a trait predicted to evolve under adaptation to high-density conditions (MacArthur and Wilson, 1967; Mueller, 1988a). This reduction gave the first indication that efficency of food acquisition versus utilization might trade off in optimising larval competitive ability: he *K*-population larvae were possibly sacrificing efficiency for a greater increase in feeding rate (Mueller, 1990, 1991).

This study was later repeated with greater rigour and gave rise to similar results once more (Guo *et al*., 1991; Mueller *et al*., 1991). Subsequently, another selection study sought to replicate these results using only larval crowding. The CU set of populations were maintained at high larval density, in contrast to their low larval density controls, named UU (Mueller *et al*., 1993; Joshi and Mueller, 1996). Additionally, their geographical origin also differed from that of the *r*- and *K*-populations. Similar to the *K*-populations, the CU populations also evolved increased larval feeding rate and foraging path length, compared to the UU (Joshi and Mueller, 1996; Santos *et al*., 1997; Sokolowski *et al*., 1997), and they showed a reduction in food to biomass conversion efficiency as well (Joshi and Mueller, 1996). Additionally, the CU populations evolved increased larval tolerance to urea and ammonia (Shiotsugu *et al*., 1997; Borash *et al*., 1998) – both known to be components of larval metabolic waste, which could build up in crowded larval cultures (Botella *et al*., 1985; Borash *et al*., 1998).

Thus, three selection experiments yielded consistent patterns of adaptation to high larval density conditions, leading to the canonical view (Mueller, 1997, 2009; Joshi *et al*., 2001; Prasad and Joshi, 2003; Mueller *et al*., 2005; Mueller and Cabral, 2012; Mueller and Barter, 2015; Bitner *et al*., 2021), which predicted that laboratory populations of *D*. *melanogaster* selected for adaptation to larval crowding would evolve increased larval competitive ability via:

1. Increased larval feeding rate.
2. Increased tolerance to metabolic waste products.
3. Increased larval foraging path length.

This adaptation would come at the cost of reduced food to biomass conversion efficiency. This view was reinforced by observations from other studies. *Drosophila* populations selected for increased larval feeding rate also evolved greater larval competitive ability (Burnet *et al*., 1977). Another study demonstrated that *Drosophila* larvae evolved decreased competitive ability as well as decreased feeding rate as correlates of the evolution of increased parasitoid resistance (Fellowes *et al*., 1998, 1999). Selection for rapid development, with or without concurrent selection for early reproduction, in populations descended from UU also led to reduced larval competitive ability along with a reduction in larval feeding rate (Prasad *et al*., 2001; Shakarad *et al*., 2005; Rajamani *et al*., 2006).

This canonical view was subsequently challenged by three selection studies. The first two selection studies involved selection for adaptation to larval crowding in populations of *D*. *ananassae* and *D. nasuta nasuta*, respectively (Nagarajan *et al*., 2016). The third study, of which the current study is a continuation, involved *D. melanogaster* populations named MCU (Sarangi *et al*., 2016). These were descended from the UU populations. All three studies involved selection for larval crowding at slightly different egg and food combinations compared to the *K*-populations or the CU populations. The results from all these three experiments were inconsistent with the canonical view, but consonant with one another (Nagarajan *et al*., 2016; Sarangi *et al*., 2016). The larval-crowding-adapted populations evolved greater larval competitive ability via a very different set of traits than those seen to evolve in the earlier studies. Compared to their low-density controls, they evolved reduced pre-adult development time and increased time efficiency of food to biomass conversion. They did not evolve greater larval feeding rate. Nor did they evolve greater urea tolerance.

It was clear that exactly how the crowding was imposed could affect the evolutionary trajectory of a population experiencing chronic larval crowding (Sarangi, 2018). More recently, our research group has carried out a long-term selection experiment with three different sets of larval-crowding-adapted populations (Sarangi, 2018). Each set is adapted to different crowding regimes. The details of these populations are as follows:

1. MCU – reared at high larval densities similar to those of *D. ananassae* and *D. n. nasuta* (Nagarajan *et al*., 2016).
2. CCU – reared at the same larval density and vial dimensions as the MCU, but with double the eggs and food amount.
3. LCU – reared at high larval densities similar to the CU populations (Joshi and Mueller, 1996), in vials with narrower dimensions than MCU (see methods).

All three sets of crowding-adapted populations evolved greater larval competitive ability compared to their low-density controls, called MB (Sarangi, 2018; Venkitachalam *et al.,* 2023). When compared with MB populations, the LCU and CCU populations, but not the MCU populations, showed an increase in larval feeding rate (Sarangi, 2018). No evolution of urea or ammonia tolerance was seen in these populations (Sarangi, 2018).

We have recently shown that evolution of larger egg size is a consistent correlate of increased larval competitive ability in these populations (Venkitachalam *et al*., 2022). Although all three crowding adapted population sets evolved increased egg size, LCU had greater egg length and egg width than the other two. Only LCU populations evolved a significantly shorter egg hatching time than the low-density controls, with the MCU and CCU populations showing a hatching time shorter than MB, but not significantly so (Venkitachalam *et al*., 2022).

A shorter hatching time and greater egg size would likely confer increased competitive ability mainly through providing an initial advantage, or a head start, in the context of larval competition (Bakker, 1961, 1969; Mueller and Bitner, 2015). Thus, we can expect the larvae of the LCU populations to show the most potent head starts in competition. Even CCU and MCU can be expected to show greater larval head starts compared to the control MB populations.

In the current study, we have asked if these differences in egg size and hatching time are important for countering a head start imposed *against* the larvae of each of these populations. For this purpose, we have provided a common marked competitor population various durations of head start in competition against each focal population (MB, MCU, CCU, or LCU). This equates to an age disadvantage imposed on the respective focal populations. In the absence of artificially provided advantages, the common competitor is of roughly similar competitive ability as the low-density controls MB (Sarangi *et al*., 2016; Sarangi, 2018; Venkitachalam *et al.,* 2023).

We assessed the effects of providing competitive disadvantage in the following two ways:

1. To what extent did the magnitude of the age disadvantage affect the outcomes of competition for each focal population?
2. How much did each focal population affect the competitive outcomes of the head-start-receiving common competitor?

We made the following predictions –

1. *All crowding adapted populations would be less affected by a competitive disadvantage than the low-density controls (MB)*.

- This would be expected, given the evolution of greater larval competitive ability and head start mechanisms in MCU, CCU and LCU.
2. *The common competitor would also gain the most competitive performance when given a head start against MB*.

- Following from prediction 1, we can predict that the crowding-adapted populations cover up the growth lag soon and challenge the unabated growth of the common competitor. We do not expect to see this for MB populations.
3. *The populations which evolved the largest eggs and the fastest hatching time (LCU) would be least affected by the age disadvantage*.

- Within the crowding-adapted populations, greater head start mechanisms should be most important for closing the initial time gap between the larvae of the focal population and the common competitor.
4. *The common competitor would gain the least competitive performance when given a head start against LCU*.

- From prediction 3, LCU larvae should be at the lowest time disadvantage at emergence. Thus, we expect that they can start feeding and growing sooner, thereby challenging the growth of the common competitor before MCU or CCU.

We measured pre-adult development time as the indicator of competitive performance. It can be more sensitive to competition in *Drosophila* than pre-adult survivorship (Sang, 1949a; Ohba 1961 as cited by González-Candelas *et al*.; 1990; González-Candelas *et al*., 1990; S. Venkitachalam and A. Joshi, *unpubl. data*).

## Methods

### Populations used

We used three sets of populations undergoing selection for adaptation to larval crowding, experienced at different combinations of egg number and food amount (MCU, CCU and LCU). We also used one set of control populations (MB), undergoing maintenance at relatively low larval densities, and ancestral to all the selected populations. A population (OE) carrying an eye colour mutation was used as a common competitor.

Each set (MB, MCU, CCU and LCU) contains four replicate populations, ancestrally linked by replicate subscript. The replicate *i* of each set of selected populations is derived from the population MB-*i*, allowing us to treat replicate populations as randomised blocks in our analyses.

The maintenance regime of each population set is as follows:

- **MB 1-4:** low density ancestral controls. Reared at ∼70 eggs, ∼6 mL cornmeal-sugar-yeast food medium, in cylindrical borosilicate glass vials of 2.2-2.4 cm inner diameter and 9.5 cm height.
- **MCU 1-4:** larval crowding with lowest food amount. Reared at ∼600 eggs, ∼1.5 mL food. Same vial and food type as MB. At the time of the current study, they had undergone 218-219 generations of selection.
- **CCU 1-4:** same density as MCU, but egg number and food amount doubled. Same vial and food type as MB. Reared at ∼1200 eggs, ∼3 mL food. At the time of the current study, they had undergone 97-98 generations of selection.
- **LCU 1-4:** larval crowding with the highest food amount, meant to mimic the larval crowding protocol of the earlier used CU populations (Mueller *et al*., 1993). Reared in ∼1200 eggs, ∼6mL food, in borosilicate glass vials of ∼2 cm inner diameter and ∼9 cm height (approx. 6-dram volume). Same food type as MB. At the time of the current study, they had undergone 96-97 generations of selection.

### Common competitor

OE (Orange Eye) is a population of *D*. *melanogaster* with an eye colour mutation. The mutation arose spontaneously in one of our populations ancestral to an MB population (Sarangi, 2018). We used it as a marker in the competition experiment of the current study. We rear this population at a similar egg number, food volume and vial dimensions as the MB populations.

The ancestry and maintenance of all these populations have been described in detail earlier (Sarangi, 2018; Venkitachalam *et al*., 2022). Figure 1 shows a summarised diagram of the maintenance regime for each population set.

**Figure 1:**
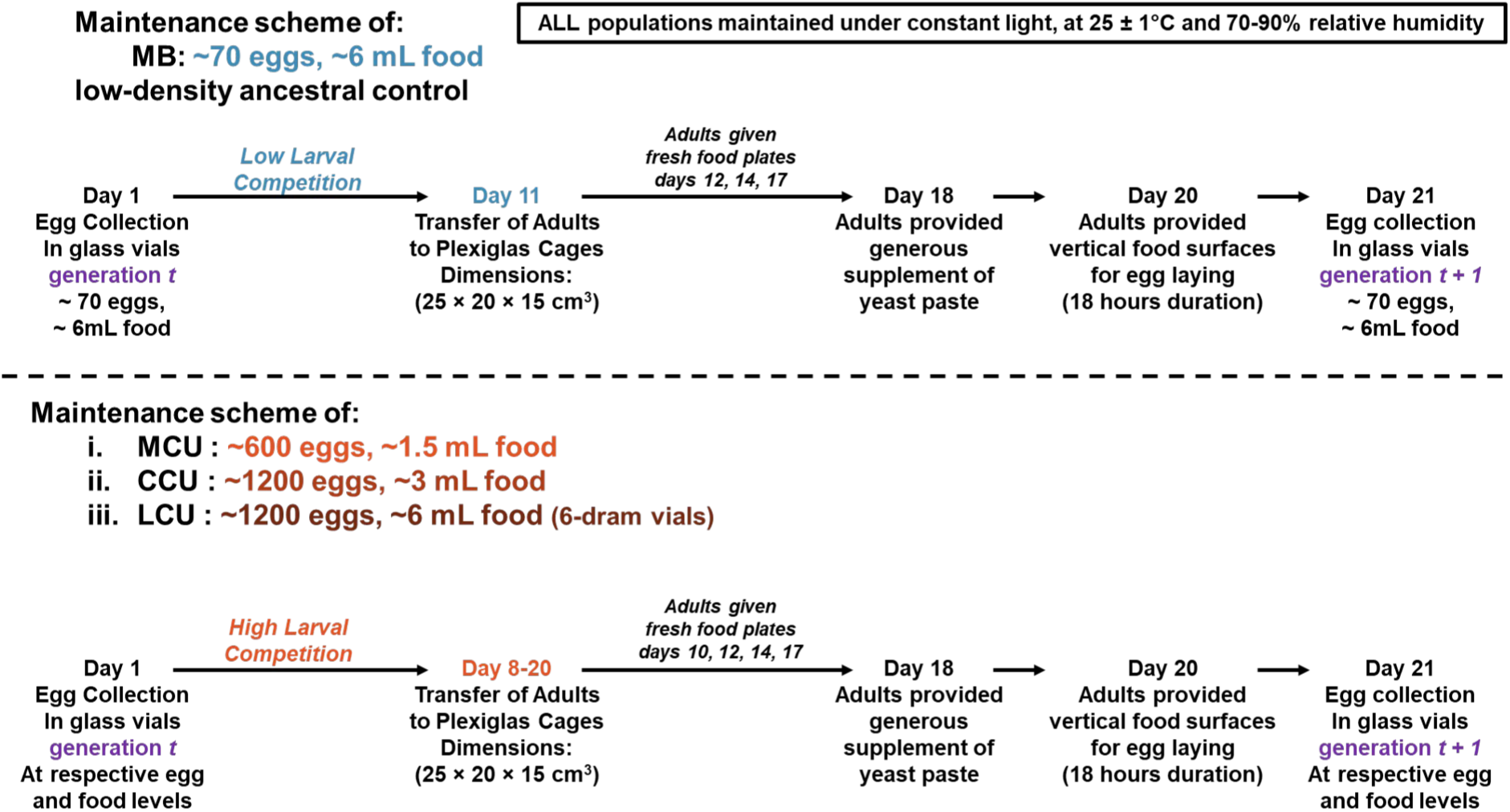
Maintenance schematic for all the populations used in the current study. In cultures with larval crowding, there is increased variation in pre-adult development time. Thus, the transfer of adults in MCU, CCU and LCU happens daily over multiple days during which adults are eclosing.

### Two generations of common background rearing

Prior to assaying pre-adult competitive ability, we standardized each population to remove non-genetic parental effects. This was done by rearing all populations in a low-density environment for two generations. At both standardization generations, we collected ∼70 eggs in ∼6 mL of food for each population, in MB-type vial dimensions. We kept a total of 40 such vials per population.

On day 11 from egg collection, we transferred the eclosed adults to Plexiglas cages. In these cages, we provided the adults with food plates smeared generously with a paste of yeast with water and a few drops of glacial acetic acid.

In the first generation of standardization, we provided a food plate for egg laying on the 13^th^ day from egg collection. On day 14, we collected eggs for the second standardisation generation.

We maintained the same protocol for generation 2 of standardisation as in generation 1 until day 13. On day 13, we first gave a regular egg-laying food plate to the adults for 1 hour, and discarded any eggs laid on it. This ensured that eggs laid from that point on would be relatively synchronized. After this step, we started the competition assay. Replicate populations 1, 3 of each set followed this schedule. For replicate populations 2 and 4 of each set, we started the experiment on day 16 <u>due to logistical reasons</u>, with the timing of yeast plate introduction also appropriately delayed.

### Assay protocol

After the 1 h egg laying window described above, we gave another set of food plates to the adults for egg laying.

These were harder agar plates, made of 2.4 g/L bacteriological agar along with yeast and sugar. The harder food surface facilitated easy removal of eggs. Once removed from the plate, we placed the eggs onto a 1 g/L transparent agar plate. We counted these eggs using soft synthetic brushes and placed them in respective vials for the assay.

The assay involved imposing larval competition in high density duo-typic cultures, i.e., focal population (one of MB, MCU, CCU or LCU) + OE, with various larval age disadvantages to the focal populations (MB, MCU, CCU or LCU). This was achieved by giving larvae of the common competitor (OE) various durations of head start (figure 2). Briefly, from the standardized populations, 200 eggs of OE were added to 2 mL cornmeal food. These eggs were added 0, 3, 5, or 7 hours before 200 eggs of a focal population (MB, MCU, LCU or CCU). Each culture thus amounted to 400 eggs in 2 mL of food. For each combination of replicate population and head start duration, we used 5 replicate vials.

**Figure 2:**
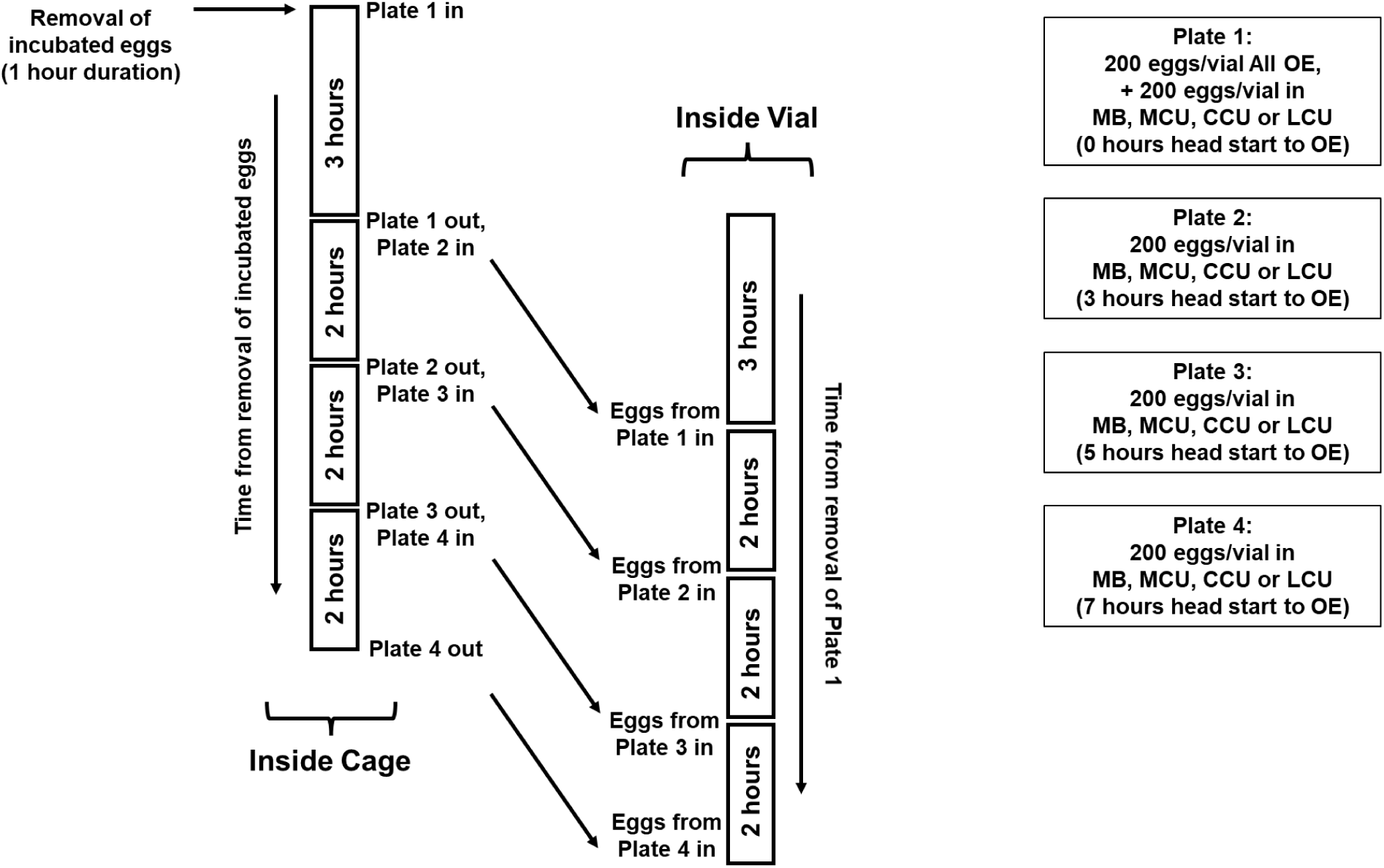
Schematic of the protocol for the pre-adult competition assay. Each plate represents a different head start duration given to OE. Plate 1 has either all the OE eggs, or eggs of the 0-hours head start treatment for focal populations. The subsequent plates contain eggs of the focal population for 3-, 5-, and 7-hours head start treatments, respectively.

We measured both pre-adult survivorship and pre-adult development time. We anaesthetised any adults eclosing from these vials (using CO_2_) and counted them. We collected and counted eclosing flies twice a day at ∼12-hour intervals for the first 3 days. This period was around day 8-10 from egg collection. Afterwards, the collection and counting frequency was relaxed to once a day. We continued this until eclosion stopped. Most flies eclosed by day 15-16 from egg collection, although a few stragglers could occasionally continue to emerge up to day 20-22. Due to logistical difficulties, eclosion checks for replicate 2 had to be stopped on the 21^st^ day from egg collection, although by this time eclosion had ceased in most vials. The development time was calculated from time of egg plate introduction until mid-point of time window of eclosion.

### Competitive performance

We wanted to test for changes in the outcome of competition, as reflected in pre-adult development time or pre-adult survivorship, due to the head start duration provided to OE. For this, we subtracted the competition outcome at 0-hours head start duration from the 3-, 5-, and 7-hour head start duration respectively. We did this for each focal population as well as for the OE competing against them.

This was calculated as follows, for both pre-adult development time and survivorship:

**a) Effect of competitive disadvantage on outcome of focal population** = Outcome with given (3, 5 or 7 h) disadvantage – Outcome without disadvantage (0 h)
**b) Effect of focal population on outcome of OE with head start** = Outcome with given (3, 5 or 7 h) head start to OE – Outcome without head start (0 h)

### Statistical Analyses

We tested the difference in competitive outcome from the 0-hour head start condition, for each population. For this, we used a mixed-model ANOVA (model III) in a fully factorial design. The blocks representing ancestry comprised a random factor. Selection and age disadvantage duration (or head start duration for OE) were treated as fixed factors. There were 3 levels of head start to OE (3, 5, 7 hours), 4 levels of selection (MB, MCU, CCU, LCU), and 4 blocks (1-4). All ANOVA were done using STATISTICA 5 (Statsoft, 1995). Tukey’s HSD was used for post-hoc pairwise comparisons. All results were plotted in R using the ggplot2 and tidyverse packages (R Core Team 2022; Wickham, 2016; Wickham *et al*., 2019).

## Results

### Effects of competitive disadvantages on focal populations (pre-adult development time)

All focal populations faced, on average, a significant increase in pre-adult development time at every level of disadvantage. This can be seen in Figure 3, where y = 0 denotes no change compared to 0 hours head start. A 3 hour age disadvantage led to a 4.6 hour increase in development time, averaged across all populations. At 5 hours of disadvantage, this value was 8.2 hours. At 7 hours, it was 10.6 hours (Figure 3). The ANOVA also revealed a significant main effect of both selection and age disadvantage for the change in development time (Table 1).

**Figure 3:**
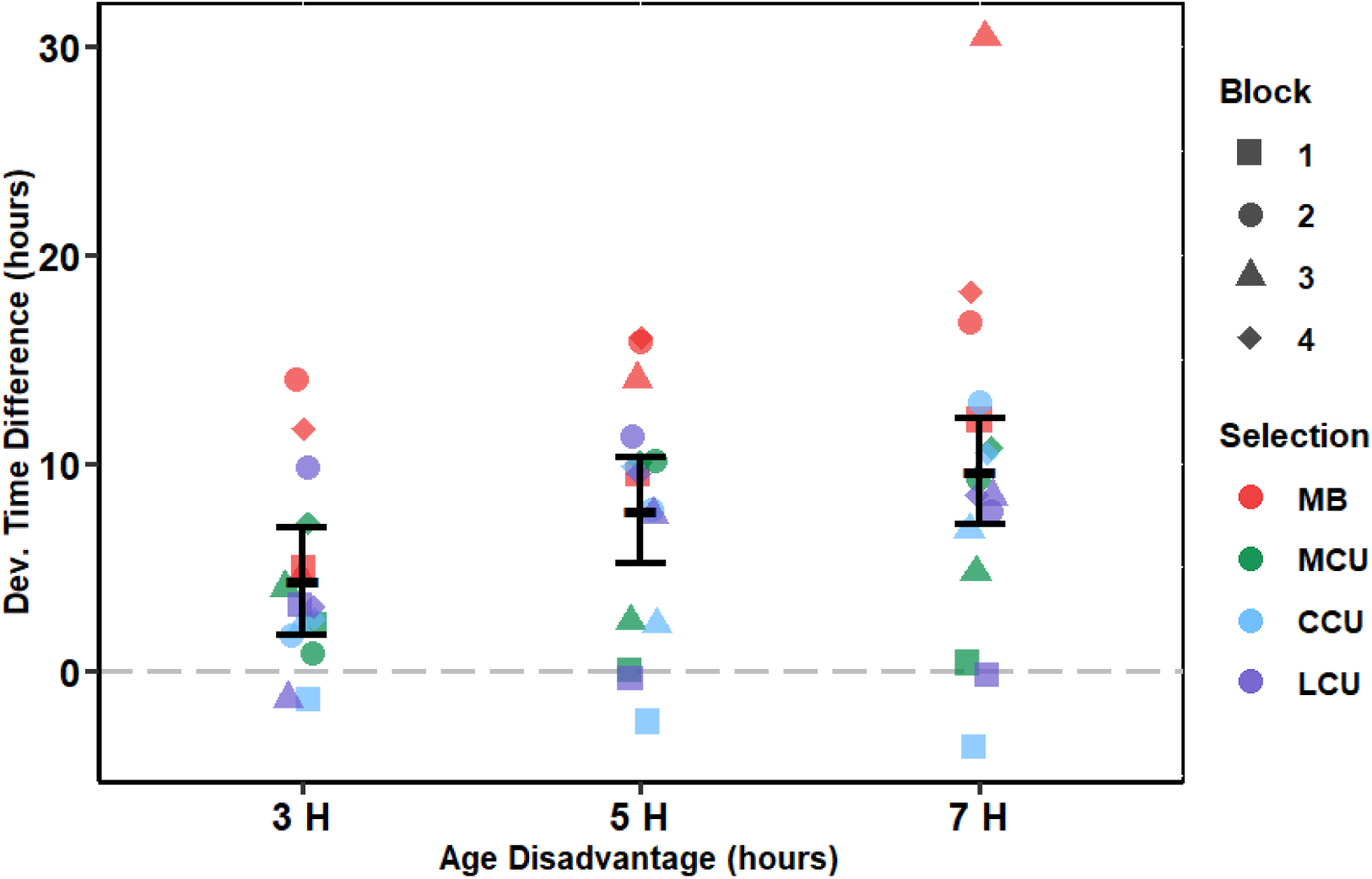
Effect of competitive age disadvantage on pre-adult development time of focal population. This is shown for the three levels of the age disadvantage given, averaged over all focal populations and blocks. The error bars show 95% confidence intervals around the mean, calculated from post-hoc Tukey’s HSD, and thus can be used for visual hypothesis testing.

**Table 1:** Mixed-model ANOVA for the effect of competitive age disadvantage on pre-adult development time of focal populations. Three factors are used – selection (fixed, four levels), age disadvantage duration (fixed, three levels), block (random, four levels). Statistically significant effects are marked. Analysis was performed on population means, therefore main effect of block and its interactions were not tested for significance.

| Effect | *df* | MS | *F* | *P* |
| --- | --- | --- | --- | --- |
| Selection | 3 | 251.602 | 34.262 | <0.001*** |
| Age Disadvantage | 2 | 114.947 | 5.200 | 0.049* |
| Selection × Age | 6 | 15.589 | 1.273 | 0.318 |

All crowding adapted populations (MCU, CCU, LCU) faced a similar increase in development time compared to the 0 hours scenario, averaged over all levels of disadvantage provided to them (Figure 4). On average, MCU development time suffered by 5.6 hours, CCU by 4.9 hours, and LCU by almost 7 hours. Each of them fared better than the MB populations, which saw an average increase of around 14 hours (Figure 4).

**Figure 4:**
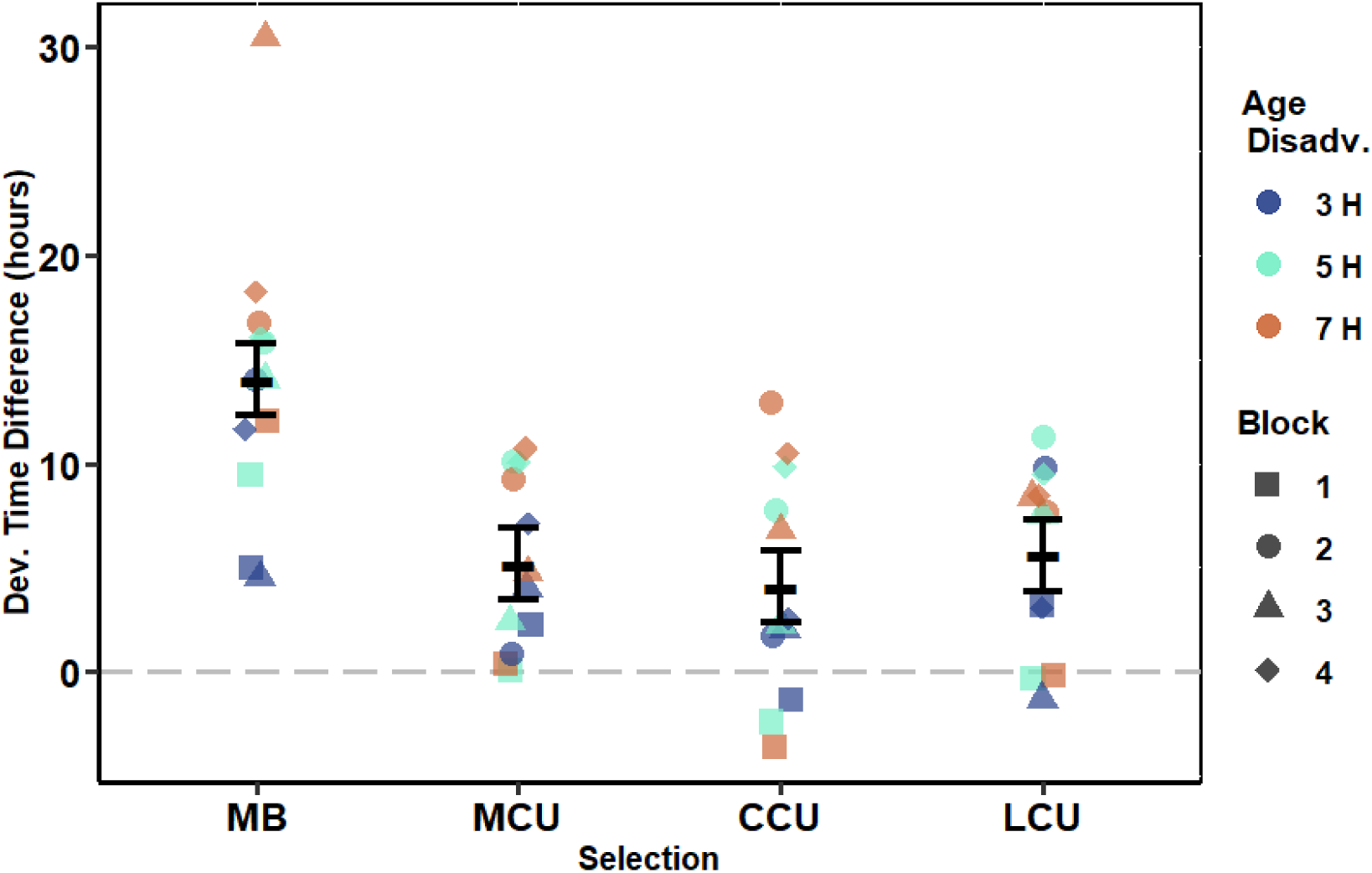
Effect of competitive disadvantage on pre-adult development time of focal population. This is shown for the four levels of the factor selection, averaged over all age disadvantage durations and blocks. The error bars show 95% confidence intervals around the population means, calculated from post-hoc Tukey’s HSD, and thus can be used for visual hypothesis testing.

The interaction of selection and age disadvantage was not significant (Table 1).

### Effects of head start duration on the common competitor against each focal population (pre-adult development time)

As seen in Table 2, the ANOVA revealed a significant main effect for head start duration. Moreover, the interaction effect between selection and head start duration was also significant. Thus, we can analyze the effect of each focal population on the competitive performance of OE at every head start duration (see Figure 5 for the following results):

- 3 hours vs. 0 hours of head start to OE: No focal population differed statistically from another in its effect on OE development time. Compared to 0 hours of head start, OE with 3 hours of head start had significantly decreased development time against only CCU (∼12 hours). On average, OE development time decreased by 7.8 and 5.5 hours when given a head start of 3 hours against MCU and LCU populations, respectively. Against MB, mean OE development time decreased by 1.4 hours.
- 5 hours vs. 0 hours of head start to OE: With 5 hours of head start, OE had significantly decreased development time in competition against every focal population, as compared to the scenario without any head start. No focal populations differed from each other in their effect on OE development time, but the pattern was that –OE development time decreased by around 24 hours in competition against both LCU and CCU, while this value was lower at 17.4 hours vs. MCU, and only 8.2 hours vs. MB.
- 7 hours vs. 0 hours head start to OE: Compared to the 0 hour head start scenario, OE had significantly shorter development time with 7 hours of head start in competition against each respective focal population. The development time decrease of OE against both CCU (35.3 hours) and LCU (38 hours) was significantly different from the development time decrease of OE against MB (10.4 hours). Moreover, the decrease of OE development time against LCU was also significantly more than the development time decrease against MCU (20.7 hours). A similar, but non-significant, pattern was seen for decrease in development time in competition vs. CCU and vs. MCU.

**Figure 5:**
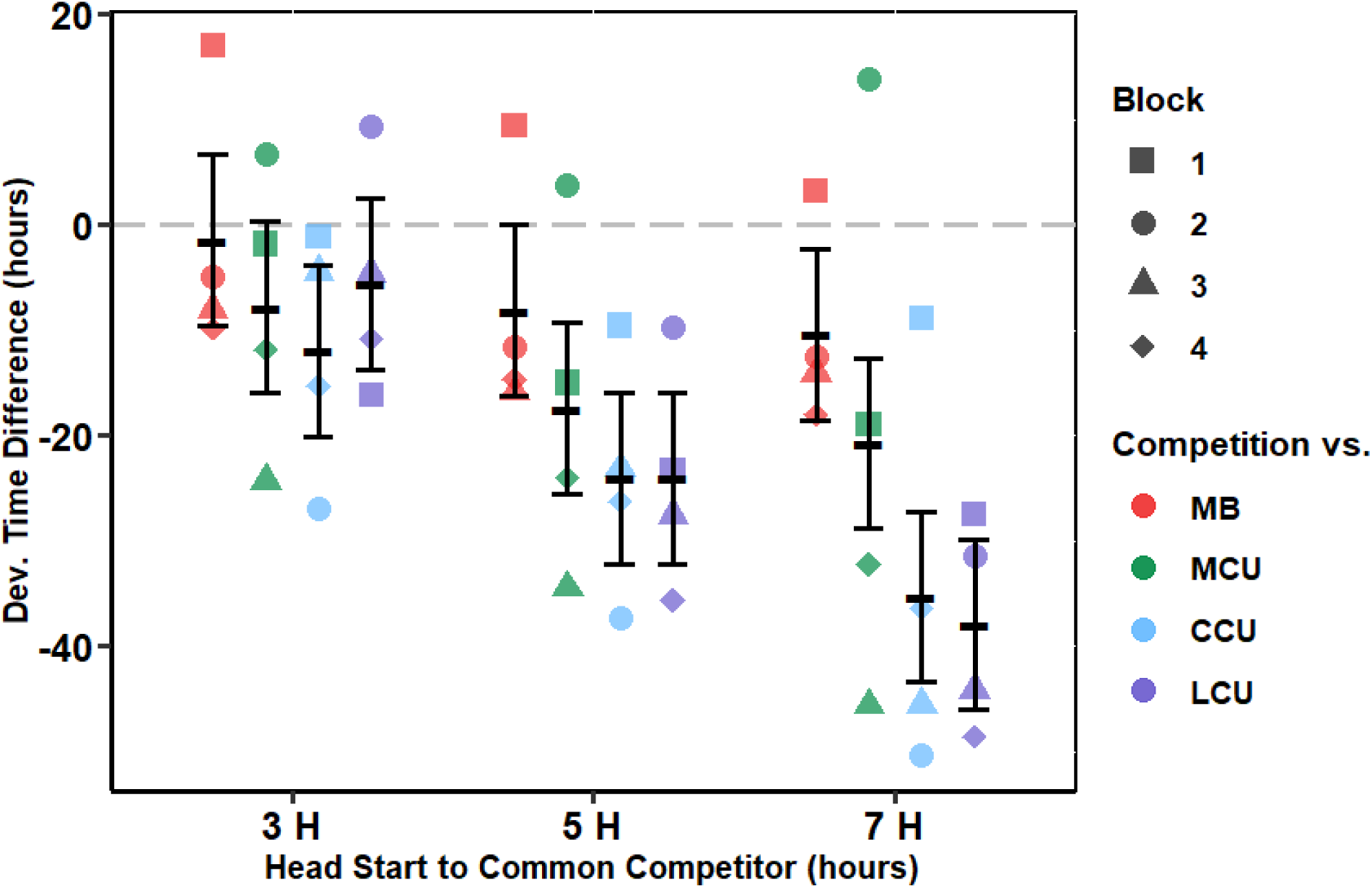
effect of focal populations on pre-adult development time of OE (common competitor) with head start. This is shown for the three levels of the age disadvantage and the four levels of focal populations, averaged over all four blocks. The error bars show 95% confidence intervals around the population means, calculated from post-hoc Tukey’s HSD, and thus can be used for visual hypothesis testing.

**Table 2:** Mixed-model ANOVA for the effect of focal populations on pre-adult development time of OE with head start. Three factors are used – selection (fixed, four levels), head start duration (fixed, three levels), block (random, four levels). Statistically significant effects are marked. Analysis was performed on population means, therefore main effect of block and its interactions were not tested for significance.

| Effect | *df* | MS | *F* | *P* |
| --- | --- | --- | --- | --- |
| Selection | 3 | 745.391 | 1.714 | 0.233 |
| Head Start Duration | 2 | 1527.034 | 36.937 | <0.001*** |
| Selection × Head Start | 6 | 113.931 | 2.990 | 0.033* |

### Pre-adult survivorship

Providing an age disadvantage did not impose any significant changes on the mean pre-adult survivorship of any focal population (Data not shown). Additionally, the common competitor did not experience any statistically detectable change in its pre-adult survivorship due to any of the head start durations provided (Data not shown).

## Discussion

### Overall patterns

We started the study with four predictions, as mentioned in the Introduction. These predictions have fared thus –

1. *All crowding adapted populations would be less affected by a competitive disadvantage than the low-density controls (MB).* This was observed. MB development time increased significantly more with age disadvantage than MCU, CCU or LCU (Figure 4).
2. *The common competitor would also gain the most competitive performance when given a head start against MB.* From Figure 5, this was rejected. MB induced the least decrease in OE development time when the latter received head starts (i.e., MB was most detrimental to OE competition) compared to MCU, CCU or LCU.
3. *The populations which evolved the largest eggs and the fastest hatching time (LCU) would be least affected by the age disadvantage.* This was rejected. No difference in the development time change was observed between any crowding-adapted population (Figure 4).
4. *The common competitor would gain the least competitive performance when given a head start against LCU.* This was rejected. The common competitor gained the **most** performance (decreased development time) against LCU, followed closely by CCU (Figure 5).

All populations selected for adaptation to larval crowding were affected to similar degrees by their competitive disadvantage (Figure 4). Moreover, the crowding-adapted population which evolved the smallest increase in head start mechanisms (MCU) reduced the competitive performance of the common competitor to the greatest extent (Figure 5).

We know from previous experiments that the larval competitive ability of OE, the common marked competitor, is lower than that of MCU, LCU and CCU. It is similar to that of MB, when tested across a range of egg and food combinations (Sarangi *et al*., 2016; Sarangi, 2018; Venkitachalam *et al.,* 2023). The current study supports this and shows that MB larvae were most badly affected by the head start to OE (Prediction 1, Figure 4).

Predictions 2, 3 and 4, however, do not hold up to empirical tests. This unexpected pattern of results might have an underlying explanation in the effectiveness and tolerance of the focal populations.

### Effectiveness and tolerance

We define *effectiveness* as the amount of growth rate inhibition a larva imposes upon its competitors. Conversely, a larva can also show some amount of *tolerance* to this growth inhibition imposed by other competitors on itself (usage inspired by Joshi and Thompson, 1995). These terms have been alternatively defined and formulated as aggressiveness and sensitivity (Breese and Hill, 1973), pressure and response (Mather and Caligari, 1983) or aggression and response (Eggleston, 1985; Hemmat and Eggleston, 1988, 1990), although in those formulations response/sensitivity is inversely related with tolerance.

We performed a subsequent experiment to measure effectiveness and tolerance of the focal populations at the same density as the current study i.e., 400 eggs in 2 mL food (Venkitachalam *et al.,* 2023). The measurements were derived from the formulae for ‘intrinsic competitive ability’ used by Joshi and Thompson (1995). For pre-adult development time, the patterns for both effectiveness and tolerance of each set of focal populations were as follows:

MCU > CCU > LCU > MB

The MCU populations showed the highest effectiveness and tolerance against OE. The MB populations showed almost no change in effectiveness or tolerance compared to OE (i.e., they had the same competitive effect on OE larvae as the OE themselves).

We can consider the increase in development time due to age disadvantage to the focal populations as a decline in tolerance. Similarly, decreasing OE development time as the duration of head start to OE increases can be considered as a decline in the effectiveness of the respective focal population that competes against OE.

For seeing change in tolerance, Figure 6 shows the least square regression of focal population development time on age disadvantage duration. Exact values of intercepts and slopes are given in Table 3. The positive slope of development time vs. age disadvantage represents decline in tolerance. For all populations, there is a general decline in tolerance with increasing age disadvantage (Figure 6). MB has the most severe decrease in tolerance as OE is given a head start against it. This also follows from prediction 1 in the introduction (i.e., MB is expected to suffer the most from an age disadvantage). As MB has the lowest tolerance even without any head start to OE (S. Venkitachalam and A. Joshi, unpubl. data), and also suffers the greatest decline in tolerance with age disadvantage, we can assume that the steepness of the decline in tolerance with age disadvantage is proportional to the tolerance measured without any age disadvantage. With this assumption, based on the measurements made on tolerance without any age disadvantage (S. Venkitachalam and A. Joshi, unpubl. data), MB should suffer the greatest decline, followed by LCU, followed by CCU, and finally by MCU, with the latter showing the lowest decline with head start to OE. However, all three crowding-adapted population sets show similar slopes (Figure 6). Thus, they suffer similar levels of decline in tolerance against OE that are given head starts against them. Based on measured tolerance, MCU should show a lower decline than CCU, with LCU showing the greatest decline among the three. The lack of a difference likely means that the greater egg size and faster hatching time of LCU and CCU help them offset a greater decline in tolerance. This provides some validation for prediction 3.

**Figure 6:**
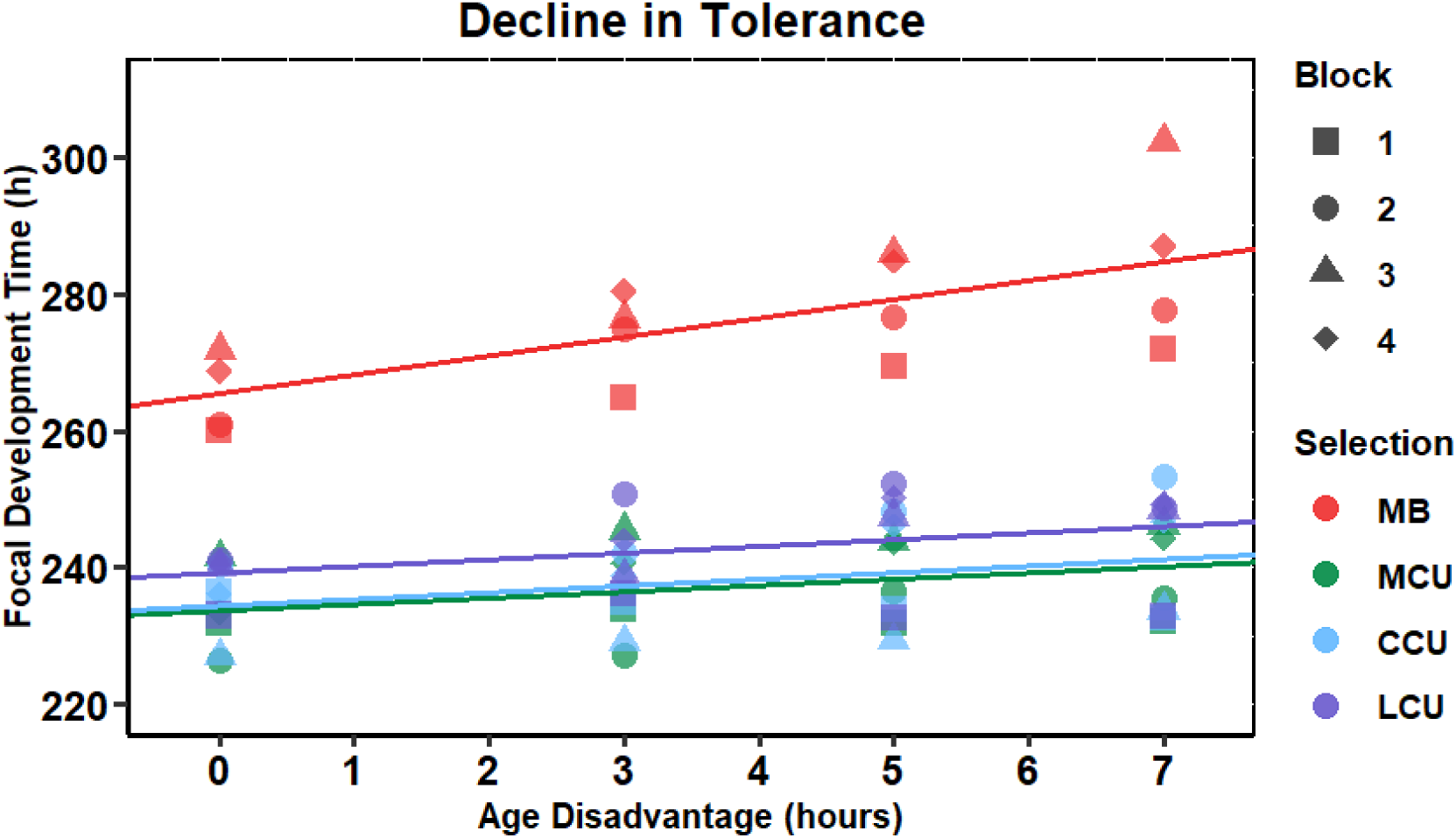
Pre-adult development time of focal populations vs. age disadvantage durations to the focal populations (by providing head start to OE). The positive slope of the least squares regression line denotes a decline in tolerance of focal populations. Exact values of slopes and intercept are given in Table 3.

**Table 3:** Intercepts and slopes for the least squares regression lines in figures 6 and 7.

|  | **Fig. 6: Tolerance Decline** |  | **Fig. 7: Effectiveness Decline** |  |
| --- | --- | --- | --- | --- |
|  | Intercepts | Slopes | Intercepts | Slopes |
| **MB** | 265.543 | 2.759 | 271.195 | -1.601 |
| **MCU** | 233.620 | 0.931 | 307.770 | -3.113 |
| **CCU** | 234.389 | 0.978 | 311.261 | -5.081 |
| **LCU** | 239.266 | 0.967 | 314.348 | -5.581 |

However, the current argument is not consonant with the inference drawn from results in figure 4. Those seem to suggest that greater head start mechanisms (egg size and hatching time) of LCU and CCU do not help offset competitive age disadvantages any better than MCU. From the perspective of tolerance decline, the lack of a difference between the crowding-adapted populations highlights a possibly nuanced rather than straightforward role larger eggs or faster hatching may be playing in the process of competition.

Figure 7 shows the least squares regression of OE development time on head start duration. Exact values of intercepts and slopes are given in Table 3. We have plotted this for competition of OE vs. each focal population. The decrease in slope of development time vs. head start shows the decline in effectiveness of the focal populations. For the crowding-adapted populations, increasing head start durations lead to different declines in effectiveness (Figure 7). The rates of decline largely follow the patterns seen for effectiveness measurements without any head start to OE (S. Venkitachalam and A. Joshi, unpubl. data). MCU, which has the greatest effectiveness, shows the lowest decline on increasing head start duration to OE. This is followed by CCU and LCU, with the latter having a marginally lower slope, i.e., greater decline in effectiveness. Even though CCU and LCU suffer more than MCU in effectiveness decline, we cannot preclude the possibility that greater egg size and faster hatching time in these populations may be preventing an otherwise even steeper decline against increasing age disadvantage. Thus, prediction 4 is left unresolved.

**Figure 7:**
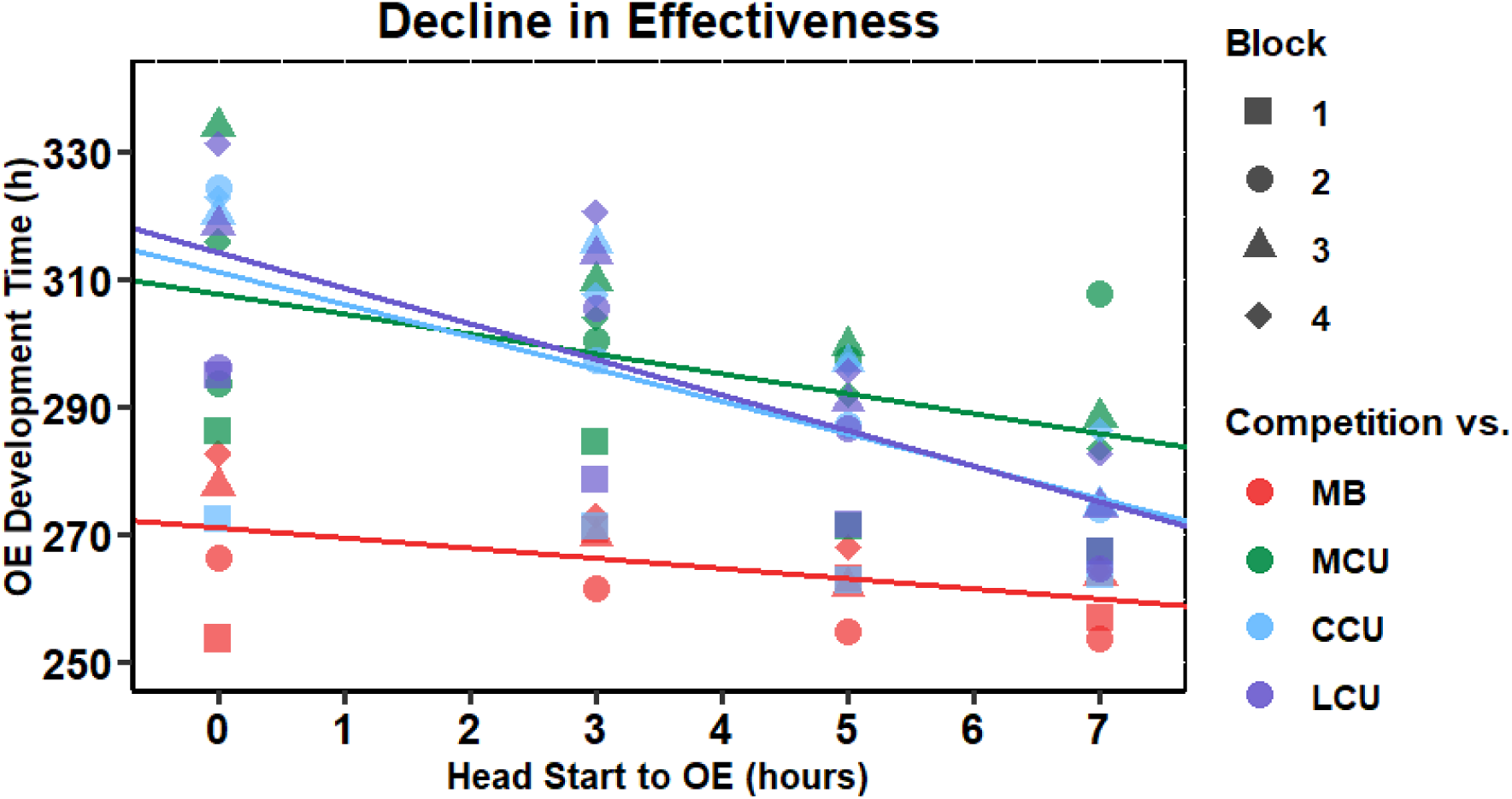
Pre-adult development time of OE against each focal population vs. head start duration to OE. The negative slope of OE development time with increasing head start duration denotes a decline in the effectiveness of focal populations. Exact values of slopes and intercepts are given in Table 3.

The major exceptions to the patterns seen for effectiveness measurements are the MB. These low-density controls have the lowest measured effectiveness in competition with OE without any head start (Venkitachalam*et al*., 2023) and yet display the least decline in effectiveness with head starts given to OE (Figure 7). This is also in contrast to the observed patterns of tolerance decline (Figure 6). We have provided a possible explanation for this in the extrapolations section below.

### Extrapolations

Based on the trends of decline in effectiveness (Figure 7), we can calculate the head start duration to OE at which each crowding-adapted population should have zero effectiveness, on average. At this extrapolated head start duration, mean OE pre-adult development time in competition against 200 MCU, CCU, or LCU larvae, respectively, would approximate competition against 200 OE larvae. For this purpose, we assume that there remains a similar slope of decline in effectiveness of each crowding-adapted population (Table 3, Figure 7), as head start to OE increases beyond the bounds of the current study. From measurements of the subsequent experiment, we have calculated the effectiveness of MB as nearly zero against OE (Venkitachalam *et al.,* 2023). In the current study, we can simplify and assume the MB condition with no age disadvantage as the reference point for zero effectiveness. This zero point yields a development time of 271.2 hours for OE (Figure 7). With this, we extrapolate, from figure 7, the following head start durations to OE required to get zero effectiveness of the respective focal population:

vs. MB: 0 hours

vs. MCU: 11.75 hours

vs. CCU: 7.89 hours

vs. LCU: 7.73 hours

Both CCU and LCU have durations to reach zero effectiveness that nearly fall within the bounds of the current experiment. Their effectiveness is almost indistinguishable from that of OE (i.e., zero) at almost 8 hours of head start. MCU populations, owing to their high effectiveness and low slope of decline, are extrapolated to suffer nearly 12 hours of age disadvantage in order to get the same effectiveness as OE. We also note that at this zero effectiveness point, the mean pre-adult development time of each crowding-adapted population set would still be faster than OE by about 25-30 hours.

Further head starts to OE from this extrapolated duration may yield further loss in effectiveness, going below zero. This is observed in the MB populations, as their effectiveness declines to negative values with increasing age disadvantage (Figure 7). However, the MB populations already possess very nearly zero effectiveness in the absence of any imposed handicap. Given further age disadvantage, it is likely that the change in development time with head start duration may dampen somewhat as effectiveness becomes negative. At values below zero, MB would probably offer very mild competitive effects to OE. With greater head starts, there may not be much scope for effectiveness to reduce further. This may explain the somewhat strange results seen for Prediction 2 (Figure 5) and the minor decline of MB in effectiveness (Figure 7). The empirical verification of this argument will require future experiments with a larger range of head start durations to OE to test the limits of decline possible for effectiveness. Additionally, providing head starts to MB against OE should give the former positive effectiveness. This would allow the observation of changes in decline of effectiveness from a positive to negative range in MB populations.

### Other outcomes of competition

At the density tested, we saw no differences in mean pre-adult survivorship. Conducting this experiment at a higher level of crowding may cause differences in survivorship. However, this may obscure trends in pre-adult development time due to very few surviving adults in some cases (Venkitachalam *et al.,* 2023). As seen in the current study, development time is sensitive to minor changes in the dynamics of competition.

Another sensitive output of competition is adult dry weight at eclosion (Sang, 1949a; Bakker, 1961, 1969; Ohba 1961 as cited by González-Candelas *et al*.; 1990; González-Candelas *et al*., 1990; Venkitachalam *et al.,* 2023). Bakker (1961, 1969) saw changes in dry weight upon implementing multiple durations of head starts on two competing *D*. *melanogaster* populations. Nicholson (1948, as cited in Nicholson, 1955) found that upon crowding the larvae of blow flies *Lucilia cuprina*, the variation in size of eclosing adults was accentuated due to batches of larvae getting various durations of head starts. Bryant (1971) also saw changes in dry weight of emerging house flies *Musca domestica* on conferring different durations of head starts to larvae of two strains. Due to logistical limitations, our study did not involve the measurement of adult dry weight. It does, however, have potential to show differences in future studies. We know that LCU-like crowded cultures show a large amount of variation in dry weight of eclosing flies (Sarangi, 2018). Adults eclosing earlier are larger. Later eclosing flies tend to be smaller, although the last eclosions are of medium-sized flies (Sarangi, 2018). Several other studies have also mentioned or shown trends for earlier eclosing flies being larger than the later ones (Sang, 1949a; Bakker, 1969; Hughes, 1980; Venkitachalam *et al.,* 2023). Since there are large development time changes in our study, we might see differences in dry weight variation of both focal populations and OE at different head start durations.

### Mechanistic explorations

The mechanism underlying the patterns seen in our experiment may be related to the dynamics of metabolic waste build-up in a crowded *Drosophila* culture. Metabolic waste products such as urea, ammonia and uric acid are known to cause mortality or increase development time in *Drosophila* larvae (Botella *et al*., 1985). Food ‘conditioned’ by larvae of different strains can also cause differential mortality in larvae introduced later in the culture of *D. melanogaster* (Weisbrot, 1966; Dawood and Strickberger, 1969; but see Budnik and Brncic, 1975 for an exception) and related species (Budnik and Brncic, 1974, 1975). Moreover, the CU populations, which are ancestrally related to the populations used in our study, evolved different traits along the eclosion time axis (Borash *et al*., 1998). From the CU-like crowded culture, offspring of earlier eclosing flies (first 72 h of eclosion) showed greater feeding rate. Those of later eclosing flies (post 500 h of eclosion, 48-72 h window) showed greater urea and ammonia tolerance (Borash *et al*., 1998). The same study also showed that waste (ammonia) accumulation occurred steadily over 20 days from egg collection.

Thus, waste dynamics likely play a larger role in the later stages of a crowded larval competition culture. The more time a larva spends in a culture, the greater the waste it likely is exposed to, and the longer it may take to develop. This means that larvae given age disadvantages may take a longer time to develop, as seen in our results (Figures 3, 4). Larvae given a head start likely face less waste and therefore may be able to develop faster (Figure 5). The delayed role of waste may also explain why LCU and CCU populations, with greater head start mechanisms, don’t perform better than MCU. Even if the LCU or CCU larvae gain advantages early on compared to MCU in competition, the larvae of the latter could close the development time gap with potentially greater larval growth rates. Characterization of larval waste dynamics in future competition experiments with varying head starts could be used to test these predictions.

### On the importance of head starts in competition

We can potentially use results from the current study to ask another kind of question: how important are head starts in determining the outcome of competition?

The literature on plant competition is rife with studies attempting to address this question (reviewed in Ross and Harper, 1972; Wilson, 1988). Initial advantages to a plant in competition could lead to cumulative benefits, either through larger seed size (Black, 1958) or faster seedling emergence time (Ross and Harper, 1972). Ultimately, a small head start to a plant against somewhat equal competitors could provide an overwhelming supremacy to it. Newman (1973) termed this a “snowball” effect. In contrast, some studies showed that such cumulative snowballing may be limited to competition for light – for nutrient based, root competition, there was little evidence that initial advantages were supremely important (Newman, 1973; Newbery and Newman, 1978; Wilson, 1988).

Due to large differences in the biology of the model systems, such arguments from plant competition cannot directly be applied to the *Drosophila* larval competition context. However, the concept of a cumulative “snowball” effect in competition should be general enough to be explored in most model systems. Indeed, Bakker (1961) has reviewed evidence for and commented on the paramount importance of head starts in larval competition in nature.

Potentially cumulative advantages have been discussed in several studies which have looked at the ecological implications of head starts in larval competition. Populations have been seen to switch larval competitive superiority after a few hours of head start in *Drosophila melanogaster* (Bakker, 1961, 1969), and house flies *Musca domestica* (Bryant, 1971). Several experimenters have studied competition by providing larval batches different durations of head start in *Drosophila* (Sang, 1949b; Seaton and Antonovics, 1967; Gilpin, 1974). Mueller and Bitner (2015) suggested that *Drosophila sechellia* larvae on the *Morinda* fruit may obtain a survival advantage from a large head start against otherwise competitively superior larvae from other *Drosophila* species. This could occur due to the former’s significantly faster hatching time and *Morinda* toxin tolerance compared to other *Drosophila* species. Initial advantages of older larvae have been shown to be important in Tephritid fruit flies *Rhagoletis pomonella* (Averill and Prokopy, 1987), as well as different species of Ichneumonid parasitoid wasps (Fisher, 1961; Jørgensen, 2009). Based on several empirical observations, the importance of head starts through early larval hatching has been suggested for three species of Scolytid bark beetles (Beaver, 1974). Except for Bakker’s (1969) study, none of the above-mentioned studies considered the effects of head starts on populations that had evolved to become competitively superior through laboratory selection.

In our results, there does seem to be some cumulative effect along the development time axis. This is evidenced by the large decrease in development time that OE experiences after gaining only a few hours of head start (Figure 5). This is additionally supported by the relatively large increases in development time of focal populations given a small age disadvantage (Figures 3, 4). However, development time of crowding-adapted populations was not affected as badly as the MB (Figure 4). Additionally, survivorship did not change significantly across any head start duration. In the case of a “snowball” effect, we would also expect survivorship to be affected by an age disadvantage.

An additional complication arises in such arguments when effectiveness and tolerance are taken into account. We consider the simplest case of OE getting a head start against an almost equal competitor, MB. The latter’s tolerance declines steeply with increasing head start to OE (Figure 7). This indicates the existence of a cumulative snowballing effect against MB. However, a similar pattern is not seen in the case of effectiveness. As seen in figure 6, OE gets the least advantage against MB, likely due to a dampened relationship between rate of decline for negative effectiveness and head start duration. This means that OE does not get nearly as much cumulative benefit against an equal competitor as it can get against potentially superior competitors such as LCU (Figure 6). Clearly, in *D*. *melanogaster* larval competition, cumulative effects from initial advantages are not obvious – they are dampened or exacerbated by the overall competitive ability, as well as relative effectiveness and tolerance, of the competitors. Further tests with different populations, varied densities and head start durations may help uncover these nuances in the process of larval competition.

## Conclusions

We have shown that increased larval competitive ability evolved through three different chronic larval crowding regimes can help larvae of the respective populations fare better against age disadvantages in competition. However, among superior competitors, those that have evolved the greatest head start mechanisms may not show better performance against age disadvantage. Studying the effectiveness and tolerance of the competitors can highlight the subtle effects head start mechanisms may nevertheless have on the outcomes of competition. Even the effectiveness relationship with head start duration may change depending on the value of effectiveness of the population. We have discussed the possibility of waste dynamics being crucial to the underlying mechanisms of the patterns seen in the current study. Such competition experiments with artificially provided head starts can also be used to explore cumulative gains from initial advantages. From the current study, we have seen that a “snowball” effect may or may not occur depending on the considerations of effectiveness and tolerance. We end with the observation, in agreement with J.H. Sang (1949b), that such competition experiments with head starts can give more nuanced understanding over and above simple larval crowding experiments.

## Acknowledgements

We thank Avani Mital, Medha Rao, Sajith V. S., Ramesh Kokile, Bhavya Pratap Singh, Rajanna N. and Muniraju for help with the experiments. S. Venkitachalam was supported by a doctoral fellowship from the Jawaharlal Nehru Centre for Advanced Scientific Research. S. Das and A. Deep were supported during their sojourn in the lab by the Jawaharlal Nehru Centre for Advanced Scientific Research’s Project Oriented Biological Education and Summer Research Fellowship programmes, respectively. This work was supported by a J. C. Bose National Fellowship from the Science and Engineering Research Board, Government of India, to A. Joshi, a Department of Biotechnology, Government of India - JNCASR project, “Life Science Research, Education & Training”, and, in part, by A. Joshi’s personal funds.

